# Robust parametric UMAP for the analysis of single-cell data

**DOI:** 10.1101/2023.11.14.567092

**Authors:** Guangzheng Zhang, Bingxian Xu

**Affiliations:** Guangming Laboratory, Shenzhen, China; Department of Molecular Biosciences, Northwestern University, Evanston, IL 60208, USA; NSF-Simons Center for Quantitative Biology, Northwestern University, Evanston, IL 60208, USA

## Abstract

The increasing throughput of single-cell technologies and the pace of data generation are enhancing the resolution at which we observe cell state transitions. The characterization and visualization of these transitions rely on the construction of a low dimensional embedding, which is usually done via non-parametric methods such as t-SNE or UMAP. However, existing approaches become more and more inefficient as the size of the data gets larger and larger. Here, we test the viability of using parametric methods for the fact that they can be trained with a small subset of the data and be applied to future data when needed. We observed that the recently developed parametric version of UMAP is generalizable and robust to dropout. Additionally, to certify the robustness of the model, we use the theoretical upper and lower bounds of the mapped coordinates in the UMAP space to regularize the training process.

## 1 Introduction

Methods that project high-dimensional data onto a lower dimensional space emerged as one of the most important facet of the analysis of single-cell RNA sequencing data. Dimensionality reduction algorithms allow the direct visualization of the high dimensional data to aid subsequent analysis such as constructing pseudotime trajectories [1, 2, 3], tracing cell lineages [4], and identifying cell types [5, 6, 7].

These algorithms can be broadly categorized into two groups, non-parametric methods that extract information from k-nearest neighbour (KNN) graphs, with t-SNE [8] and UMAP [9] being the most popular, and parametric ones that “maps” data to a lower dimensional space with complex, nonlinear functions parameterized by a neural network, constructed through the minimization of loss functions [10, 11, 12, 13].

In most cases, non-parametric methods, requiring only a few hyper-parameters as inputs, are favored partially because of the noisiness of single cell data, which hinders the generalizability of parametric methods as they can contain thousands of parameters. Finding a robust set of parameters for parametric methods is non-trivial, and the fact that they require a training step which does not translate to additional knowledge of the system further diminishes their popularity.

However, as the size of data grows, we argue that parametric methods may have an edge over non-parametric ones for two important reasons. First, the sheer size of the data can make it extremely time consuming for methods such as t-SNE or UMAP to construct KNN graphs; and second, in the situation when one wishes to compare his/hers own dataset, which is small, to an existing large dataset, he/she cannot circumvent the need to re-do the cumbersome procedure including the construction of a KNN graph, which can be particularly difficult when one’s local machine cannot even load the large dataset. Through the proper use of parametric methods, we argue that one can train a model with a fraction of the data ensemble, and map the remaining data onto the same space, saving computational time, or map one’s own data with a pre-trained model to make easy comparisons.

Another benefit of using parametric methods lies in their tractability. While one cannot know how a small perturbation to the input data will effect the final embedding when using non-parametric methods, the impact of such small perturbation can be tested with their parametric counterparts directly. Furthermore, one can even differentiate the model output with respect to the input to compute how each input impacts the final outcome.

Here, we proposed that the parametric UMAP [13] (pUMAP) is good for analyzing single cell datasets. We showed that pUMAP can perform just as well as its non-parametric counterparts. Additionally, we show can one can incorporate in the training step of pUMAP a regularizer that mathematically ensures robustness, further enhancing its ability to treat the noisy single cell data.

## 2 Methods

### 2.1 PyTorch implementation of the parametric UMAP

Parametric UMAP is implemented with custom python code. First, KNN graph is constructed using the NNDescent function from the pynndescent package in python with recommended parameters [14]. The output of NNDescent is then used to construct a fuzzy simplicial set using the fuzzy_simplicial_set function from the umap package [9] from which we obtain a graph representation of the data. We built the neural network used in all experiments in PyTorch [15] with three layers of fully connected neurons with ReLU (rectified linear unit) activation. The fuzzy set cross entropy, as defined in (12), can be expanded to be:

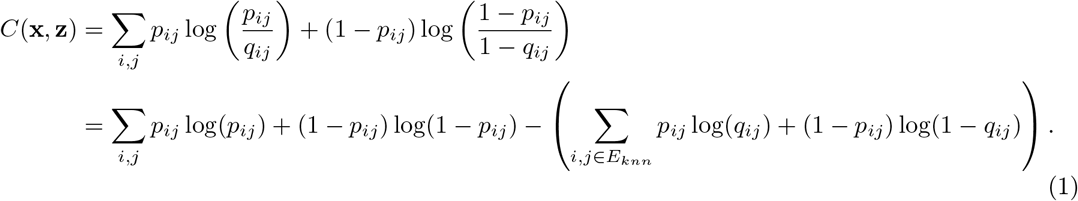

Since *p*_*ij*_ are fixed and only *q*_*ij*_ would change with the parameters of the neural network, the loss function is therefore:

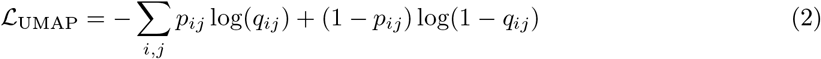

In practise, we do not sum over all pairs of points. Instead, we first sample from the set of edges to generate an edge list. Because the graph is sparse, we can assume that a randomly chosen pair of nodes will not be connected. As a result, we generate a list of negative samples by permuting the previously generated edge list. With this setup, the loss function becomes:

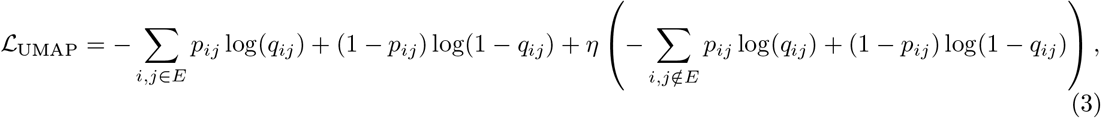

where *η* is the negative sample strength. In our setup, we call pUMAP trained with this loss the probability-based implementation as *p*_*ij*_ represents probabilities. This loss function can be simplified further into a graph-based implementation by setting *p*_*ij*_ to one if it is greater than zero:

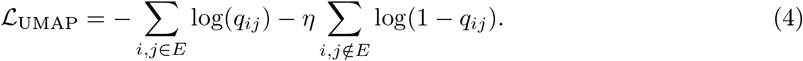

For each edge, we generated five negative samples by default. The network is trained with the ADAM optimizer [16] with a learning rate of 0.001.

### 2.2 Single cell datasets

The pancreatic development dataset [17, 18] is downloaded from: https://figshare.com/articles/dataset/Pancreas_development/13078955?backTo=/collections/CellRank_for_directed_single-cell_fate_mapping_-_datasets/5172299.

The hippocampal development dataset [19, 20] is downloaded from: http://pklab.med.harvard.edu/velocyto/DentateGyrus

### 2.3 Data preprocessing

Downloaded data were preprocessed using scanpy [21] in python. Top 300 highly variable genes were first selected with `scanpy.pp.highly_variable_genes` with flavor `seurat_v3` [22]. Data were then normalized to have the same library size with `scanpy.pp.normalize_total` to a target sum of 10000 and scaled with `scanpy.pp.scale`. Z-scores of highly variable genes were then used as input to the neural network.

### 2.4 Regularizing pUMAP

The theoretical upper and lower bound were computed using the CROWN method [23] (see SI for reproduced detail) using the auto_LiRPA package [24, 25, 26] with *ϵ* = 0.2.

For each cell *i*, we obtain an upper and lower bound for each neural network output (two in this case), 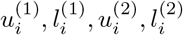, where *u* and *l* denotes the upper and lower bound respectively and the superscript (1) denotes the UMAP dimension. From these upper and lower bounds, we construct the bound loss as:

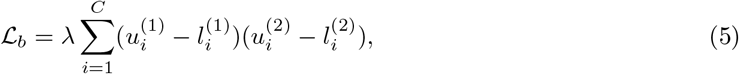

where *λ* is a scaling term that takes into account of the size of the UMAP space. This loss can be generalize to a higher dimensional UMAP space by:

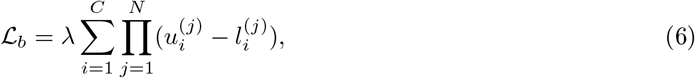

where *N* is the dimensionality of the UMAP space. During training, we incorporated ℒ_*b*_ at the very end of the training procedure. To ensure that we only constrain the bound of the output but do not change the shape of the embedding, we added a reconstruction loss:

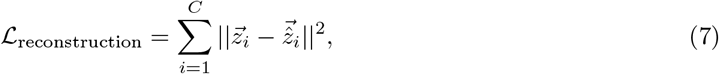

Where 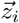 is the output of the network without the constraint imposed by ℒ_*b*_ and 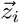 is the output of the network being trained. Hence, the loss function used during the regularization step consists of three terms in total:

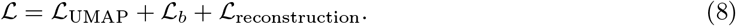

## 3 Results

### 3.1 Parametric UMAP

pUMAP [13] uses neural networks to parameterize complex functions that map gene expression data onto a lower dimensional space of an arbitrary dimension (Figure 1A). Similar to the original UMAP [9], pUMAP begins with the construction of a k-nearest neighbour (KNN) graph from the high dimensional space and computes a weight for the edge that points to *j* from *i* that scales with their local distance:

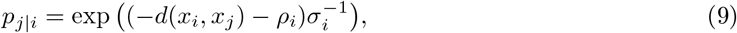

where *ρ*_*i*_ is the distance between *x*_*i*_ with respect to its nearest neighbour. Next, a symmetrized edge weight between point *i* and *j* is computed as:

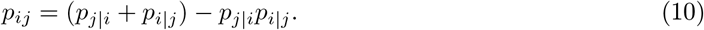

**Figure 1:**
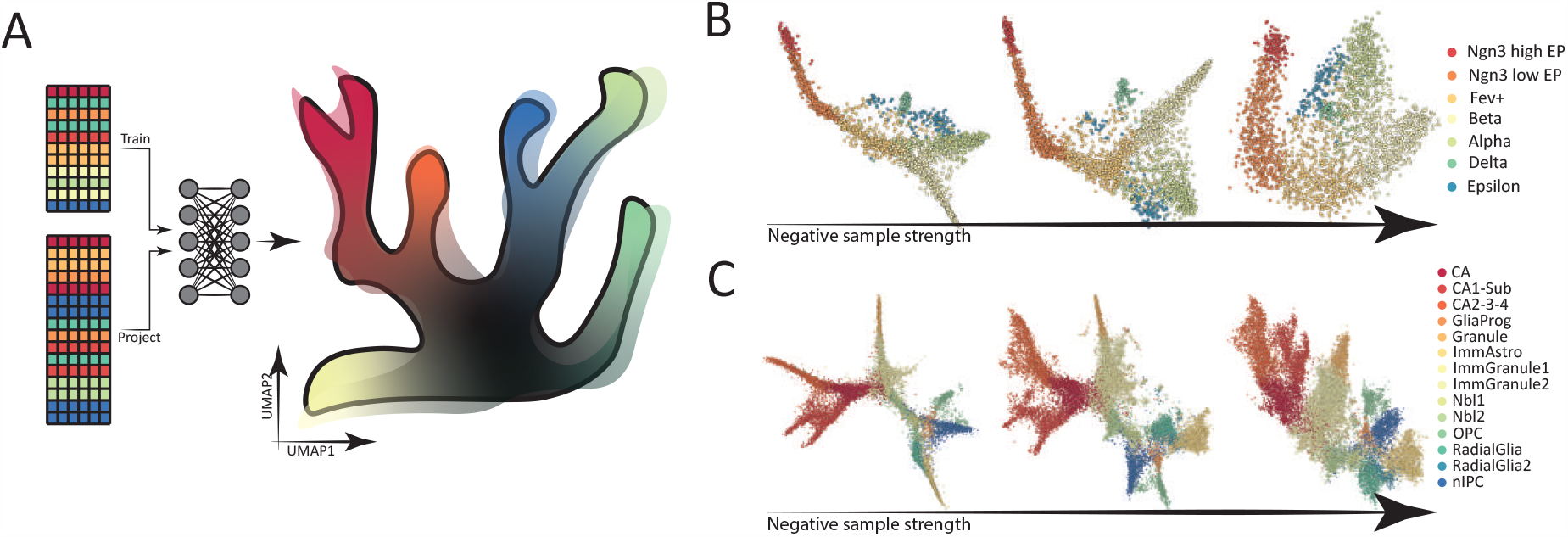
A: pUMAP is capable of efficiently projecting future data onto the same space as the training data. The effect of negative sample strength on the overall structure of the low dimensional embedding produced by trained pUMAP for pancreatic (B) and hippocampal (C) development.

For the low dimensional space, “distance” between embedded points *z*_*i*_ and *z*_*j*_ is quantified by:

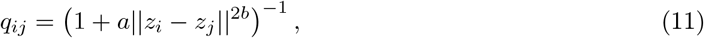

where *a* and *b* are hyperparameters to be selected. Lastly, a suitable embedding is then constructed by optimizing (see details in SI) the fuzzy set cross entropy, defined as:

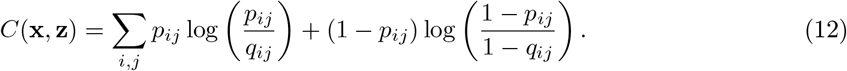

### 3.2 Parametric UMAP separates known cell types

To see whether pUMAP is suitable for analyzing single cell data, we first investigated if it can map gene expression onto lower dimensions that separates annotated cell types. We ran our own implementation of the pUMAP on two separate datasets. One charted the differentiation of pancreatic cells [18, 17] and the other characterized that of the hippocampus [20], both containing branching trajectories. We found that pUMAP can separate different cell types well (Figure 1B, C). Additionally, we observed that when the graph-based (See details of the graph-based and probability-based implementation of pUMAP in methods) version of the pUMAP was implemented, one can decrease negative sample strength to enhance the ability of the final embedding to discern between major branches while losing resolution within the individual branch (Figure 1B, C).

### 3.3 Parametric UMAP shows generalizability

Next, we evaluated the performance of pUMAP when trained on only a subset of the full data (Figure 2A, left). We sought to find out two things. First, is a small subset of the data sufficient to separate the different cell identities within the population. If so, will the train and test dataset be projected to the same space?

**Figure 2:**
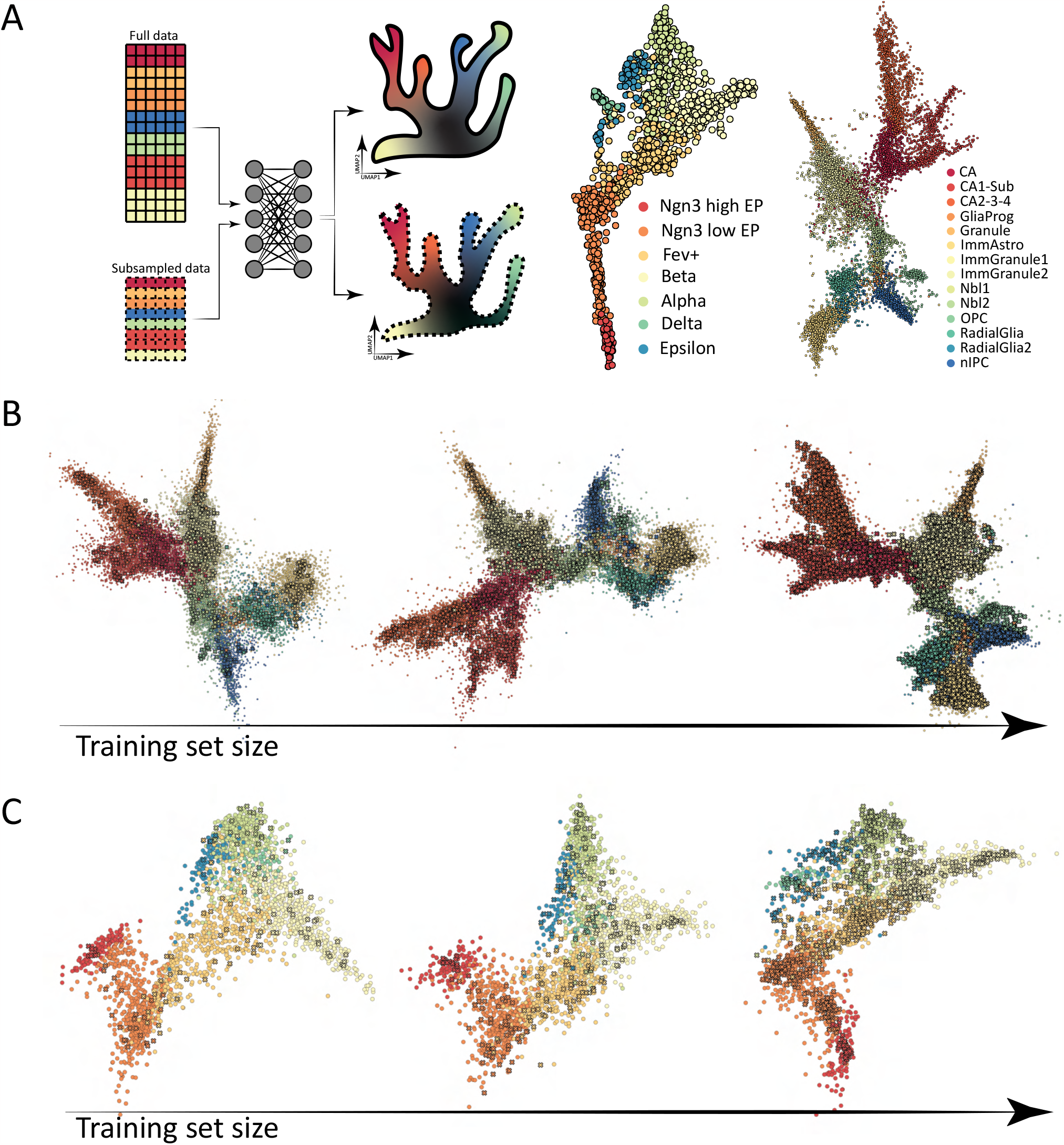
A: (right) A schematic showing a neural network trained on a subset of the full data that is later used to project the test set onto the low dimensional space. (left) pUMAP output of the pancreatic and hippocampal development trained with half of the full dataset. B and C: The effect of training set size on embedding quality for the hippocampal (B) and pancreatic (C) datasets. Neural networks were trained with 5%, 10% and 30% of the full data going from left to right. Cells used during training were plotted as crosses and the rest as dots.

By training with half of the full data, we observed that the annotated cell types are well separated for both the pancreatic and hippocampal development (Figure 2A, right), and the resulting embeddings are qualitatively similar to that obtained when the full datasets were used during the training step (Figure 1B, C).

To examine the extent to which the trained model can be generalized, we conducted training with 5%, 10% and 30% of the full data. Amazingly, we found that cell identities were well separated even when training was done with only 5% of the full data in both cases (Figure 2B,C, left plot). However, the lack of training data also resulted in a loss of resolution (Figure 2B,C), unable to recover fine structures such as the diverging development of alpha and beta cells during pancreatic development, a feature that was recovered when 10% of the pancreatic dataset was used for training (Figure 2C).

The input to our model are zscored normalized counts (see details of data preprocessing in method), which means that the input to the network for each cell can be slightly different based on the full dataset (the full data used to compute zscores) from which it is drawn and subsequently generate different embeddings. To test whether this would happen, we input zscores computed with the full datasets to models trained with zscores computed from only the training set. We observed that while zscores computed from the full and train set would be different, the full data was projected onto the same space in all test cases with train samples spreading qualitatively uniformly across the embedding (Figure 2B,C). We reasoned that as long as the train set can well-represent the full data in the sense that all cell types are present with similar proportion, as expected if the train set is a random sample drawn from the full data, the zscores for each cell would remain similar and all data will be projected onto the same space even if most are out of sample cases.

### 3.4 Parametric UMAP fills in gap in the embedding

Having shown that pUMAP can generalize when the train set can well represent the full data, we investigated the impact of having a train set that is not a good representation (Figure 3A). To do this, we first constructed a train set by removing all of the Nbl expressing cells. We hypothesized that the removal of cells that connect between the immature granules (ImmGranule1 and ImmGranule2) and radial glial cells (RadialGlia) will leave a gap region in the lower dimensional embedding (Figure 3A). As expected, we observed a clear gap region on the final embedding (Figure 3B) while the relative position of the rest of the cell types remained similar to embedding produced from models trained with the full data (Figure 1C).

**Figure 3:**
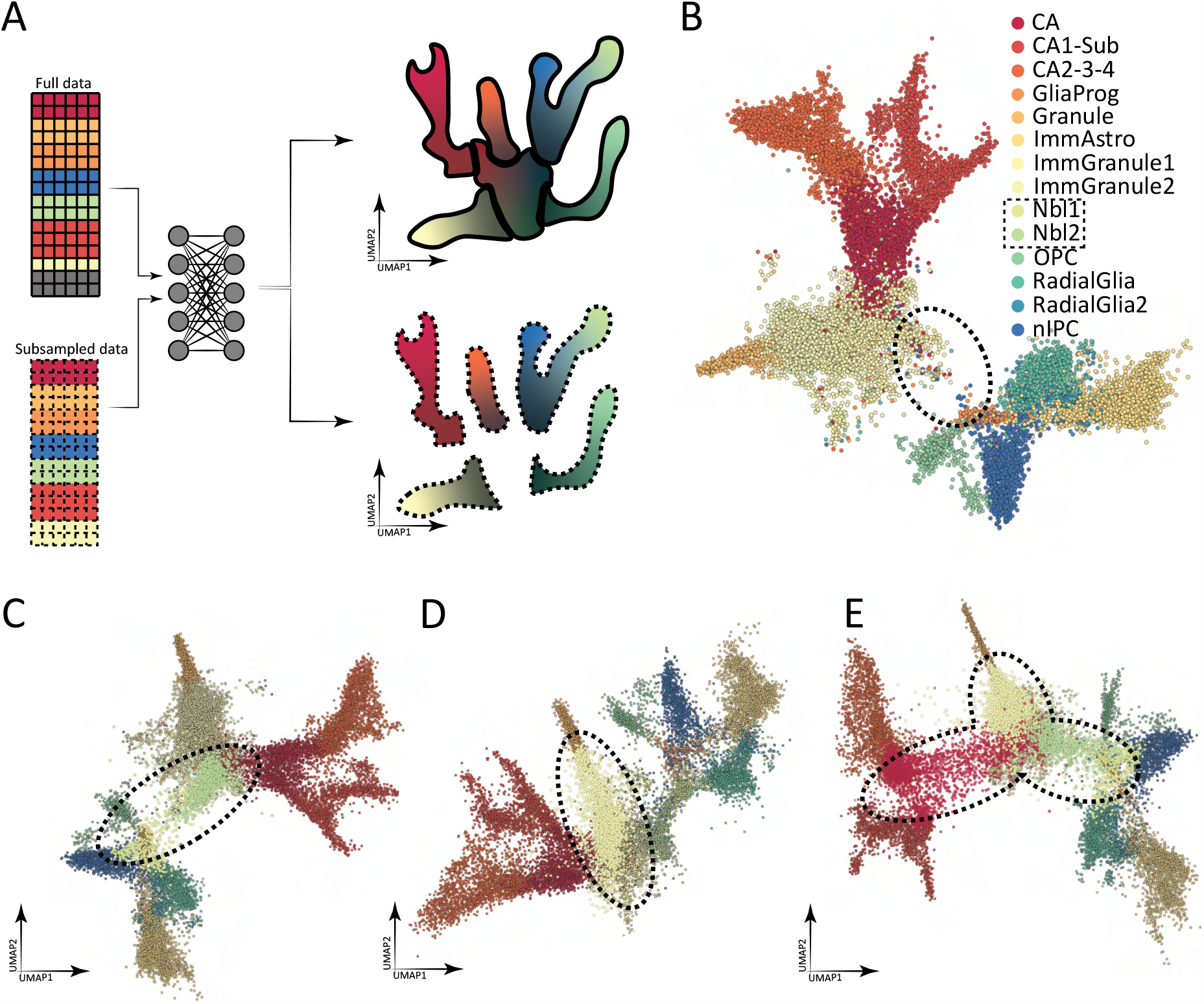
A: A schematic showing a neural network trained on the full data with just the grey cell type removed, resulting in a gap in the final embedding. B: Neural network trained on the hippocampal dataset with the Nbl expressing cells (Nbl1 and Nbl2 clusters) removed produced an embedding with a gap region. Projecting the full data using networks trained without the Nbl expressing cells (C), the immature granule cells (D), or all of the immature granule cells, Nbl expressing cells, and CAs removed (E). Data seen during training were plotted as crosses and unseen data as dots. To aid visualization, the approximate regions of the cells removed during training have been circled.

This observation led to the hypothesis that the gap regions within the embedding represent unobserved, intermediate cell types. To test this, we projected the full hippocampal dataset onto model trained without the Nbl expressing cells. Surprisingly, we found that the two clusters missing in the training set were projected to where they belong (Figure 3C). Though they do not fully cover the gap region, we have observed that the Nbl1 cluster to be located on the same side as the radial glial cells and the Nbl2 cluster on that of the immature granules, in congruence with the full model (Figure 1C) and the published embedding [19].

To further test our hypothesis that gap regions represent intermediate cell types, we conducted training with the ImmGranule2 cluster removed. This time, we observed that the model is capable of “learning” where the ImmGranule2 cluster belong, correctly projecting them so that they connect the ImmAstro cluster to the rest of the cells (Figure 3D).

Having observed that pUMAP can at least in some cases successfully fills in the gap on the low dimensional embedding with data unseen during training, we sought to test the extent of its capability by conducting training with the ImmGranule2, Nbl1, Nbl2 and CA clusters removed, resulting in at least three disconnected clusters on the low dimensional embedding. To our surprise, model trained with subsetted data can place unseen cells almost perfectly (Figure 3E), showcasing the power of using a pre-trained model.

To further confirm that pUMAP is suitable for the analysis of single cell data, we turned to test the performance of pUAMP against another major characteristic of single cell data, sparsity. To this end, we first trained a model with the full dataset and randomly set a chosen fraction of the data to zero (Figure 4A). By comparing the embedding produced with the full dataset to that from datasets that contained additional dropouts, we observed that their coordinates remain highly correlated even when 80% of the non-zero counts have been set to zero (Figure 4B). However, examination of the resulting embedding produced from data containing dropouts revealed a severe loss of resolution while cell types remained qualitatively separated (Figure 4C).

**Figure 4:**
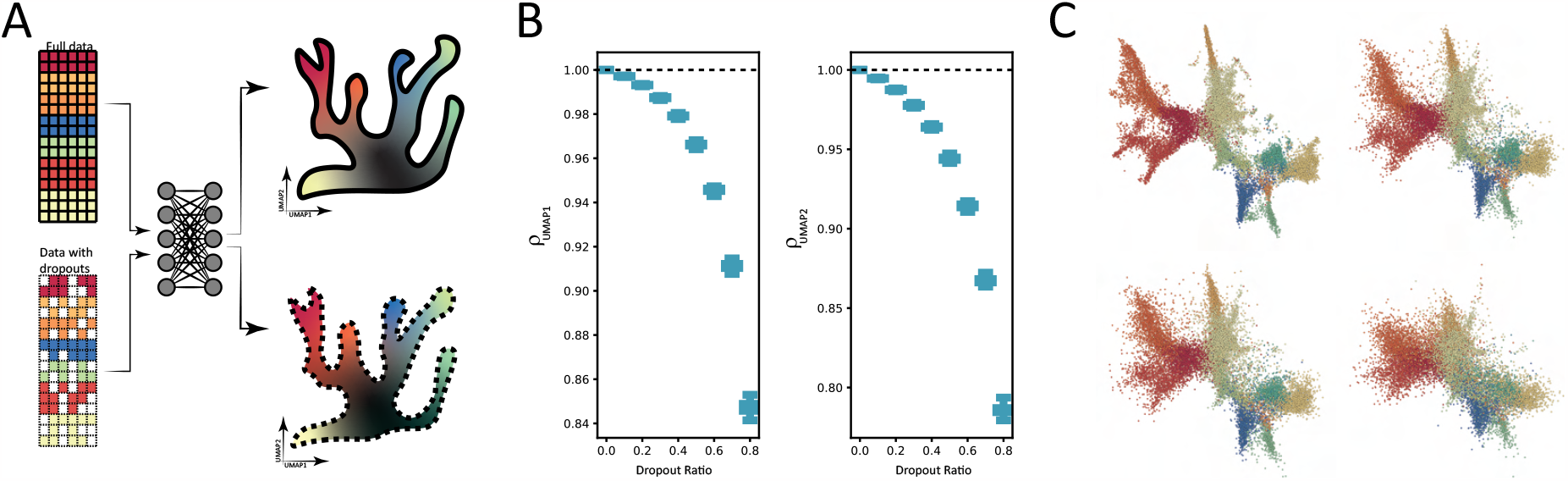
A: A schematic showing a neural network trained on the full data with induced dropouts produced an embedding similar to that trained with the full data. B: Boxplot showing the correlation coefficient between the embedding produced with models trained with dropouts and that produced with model trained without. For each dropout ratio, the same analysis were repeated 100 times. C: Example embeddings produced with a dropout ratio of 0%, 30%, 40%, and 70% respectively.

### 3.5 Robust parametric UMAP with CROWN regularization

While we have shown that pUAMP is suitable for analyzing single cell data in terms of computational cost, generalizability to unseen cell types and robustness to dropout, there still remains a vast sources of noise that could potentially be significant and diminish its usefulness. For example, recent statistical modeling of single cell data have started to consider factors such as the PCR reaction, droplet size, capture efficiency, and even ambient RNA (cell-free RNA) [27, 28]. Our inability to test the impact of these factors directly led us to take an even simpler approach of computing upper and lower bounds. In other words, for the construction of a robust model, we want to ensure that any point located near the an observation will not be projected too far away. Formally, we constraint the model by minimizing the area needed to enclose the region on the low dimensional space constructed by projecting all points on the high dimensional space that are located within a ball of radius *ϵ* centered at a data point(Figure 5A). This area can be computed analytically for neural network with simple activation functions by CROWN [23] and added to the loss function as a regularization term.

**Figure 5:**
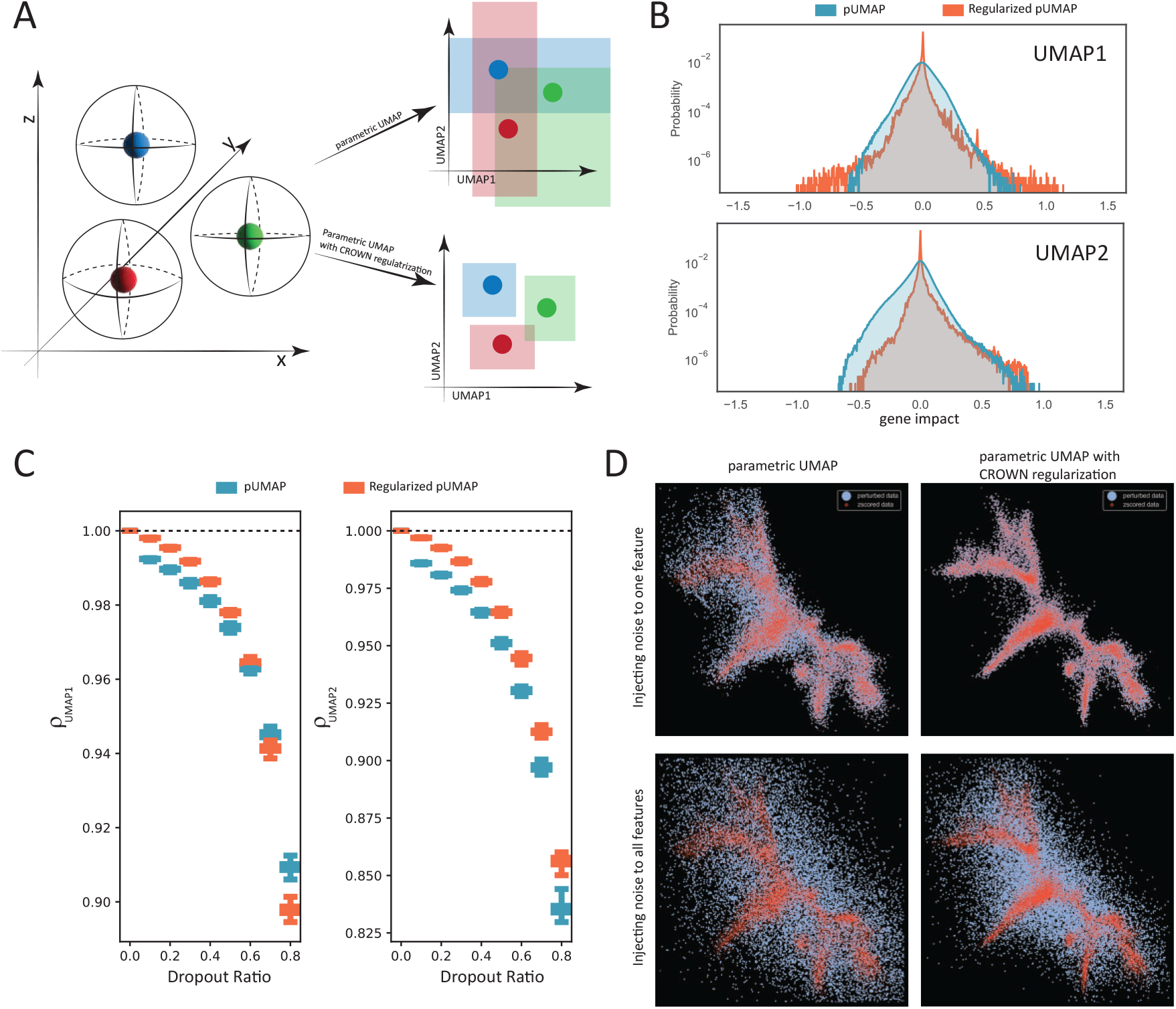
A: (left) A schematic showing a high dimensional space with three data points each being the center of a sphere with radius *ϵ* that do not overlap. (right) A schematic showing neural networks trained with and without regularization. When data are projected down to the low dimensional space using the unregularized model, their theoretical bounds showed significant intersections that can be reduced with regularization using CROWN. B: Distribution of gene impact computed from a model with and a model without regularization. Gene impact of model without regularization has a much bigger variance. C: Boxplot showing the correlation coefficient between the embedding produced with models trained with dropouts and that produced with model trained without. D: (Top row) Embedding produced from a model with and a model without regularization subjected to perturbation to one single gene. (Bottom row) Embedding produced from a model with and a model without regularization subjected to perturbation to all genes.

To test whether the addition of this regularization term increases robustness, we took advantage of the parametric nature of our approach to quantify the effect of a small change in the input by directly computing the gradient. Denote the entirety of the input data by **X** with shape *C* by *G* which represents the number of cells and genes respectively. For each gene in a particular cell (a row in **X**), we can compute how **X**_*i,j*_ impacts the output 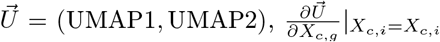, using automatic differentiation and sum over *c* to get the overall impact of this gene:

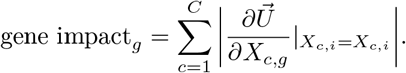

We computed the gene impact for all genes from one model that has the additional regularization term and another without. As expected, we observed that the distribution of gene impact for the regularized pUMAP has a sharp peak near zero that is much taller than that of the other pUMAP model without regularization (Figure 5B). Additionally, we found that regularized pUMAP is even more robust to dropout noise, having a higher correlation with the unperturbed embedding especially when the dropout ratio is low (Figure 5C).

Lastly, we tested the strength of regularized pUMAP by targeting specific genes. For pUMAP without regularization, we observed that perturbing the one gene with the highest gene impact can severely decrease the quality of the embedding (Figure 5D, upper left). However, when the same input was given to the regularized pUMAP, we observed little effect (Figure 5D, upper right). To ensure our observation is not an artifact of over-fitting, we contaminated all data points and observed a loss of structure in both cases (Figure 5D, bottom row).

## 4 Discussion

In this work, we showed that the parametric UMAP is a suitable tool for the analysis of single-cell RNA seq data. In a world where an enormous amounts of new data are being generated everyday, using pUMAP allows one to generate a model from a small subset of the data to reduce computational cost. Additionally, even in the situation where one wants to train a model with as much data as possible, the batch-wise training scheme adopted by pUMAP allows one to partition the data for efficient memory allocation. Moreover, we showed that in at least some cases, the underlying neural network that parameterized pUMAP appeared to have learned something about the biology, capable of filling in gaps within the embedding with cells never seen during training.

To enhance the robustness of pUMAP when it is impossible to account for the effect of all factors involved in the data generation process, we exploited the parametric nature of our approach and theoretically computed an upper and lower bound [23, 24, 25, 26], which is then incorporated as an regularization term in the loss function. We showed that regularizing with CROWN [23] greatly enhanced the robustness of the model and confirmed that our observation is not an artifact of overfitting by perturbing all genes simultaneously.

While we showed that regularizing with CROWN is a viable way to enhance the robustness of the neural network underlying pUAMP, it is not guaranteed that such approach can remove batch effect.

Future work is still needed especially in combining pUMAP with other analysis. For example, given our observation that, at least in some cases, gap regions on embeddings produced by pUMAP represent unseen cell types, one can train a VAE [29] like model that sample directly from the UMAP space to infer the gene expression profile of cells that could fill in the gap. We envision such work may be able to identify critical points of transition in diseases and pinpoint potential genes for targeted therapy.

## 5 Supplemental material

### 5.1 Certified Robustness

Below is a review of the derivation of the upper and lower bound proposed by Zhang [23], which is hopefully a bit simpler for a non-mathematical audience to follow.

Consider a neural network with *m* layers with input dimension *n*_0_ and output dimension *n*_*m*_. In the layer *m* − 1, the pre-activated input to the *i*-th neuron is 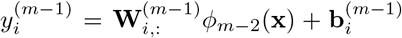. Assume we can find functions 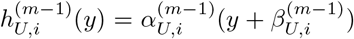 and 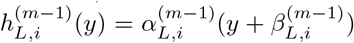 such that:

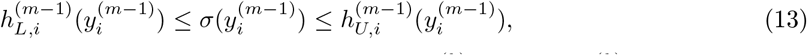

where *σ*(*y*) is some predefined activation function and *ϕ*_*k*_(**x**) = *σ*(**W**^(*k*)^*ϕ*_*k*−1_(**x**) + **b**^(*k*)^) defines the map that maps the input to the k-th layer, our goal is find functions that bound the final output of the network 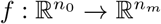. The j-th output of our network *f*_*j*_(**x**) is defined as:

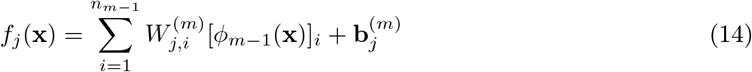

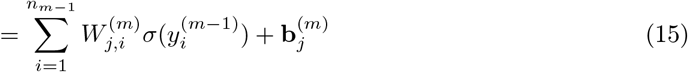

We will drop bolded letters. Employing the bound defined in (13), we have for positive 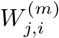:

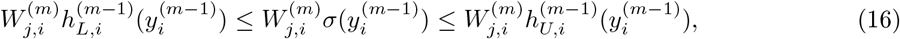

and for negative 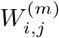:

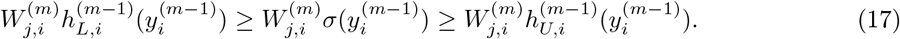

Plug (16) and (17) into (15), we can derive an upper bound, 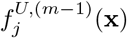 based on the upper and lower bound of *ϕ*_*m*−1_(*x*):

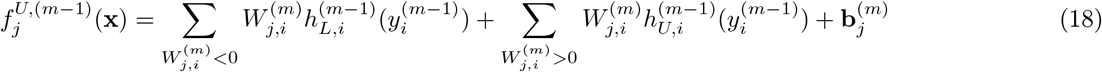

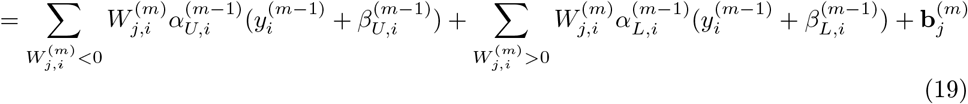

To merge these two sums, define 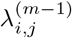 and 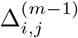:

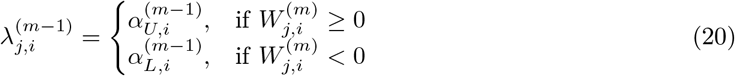

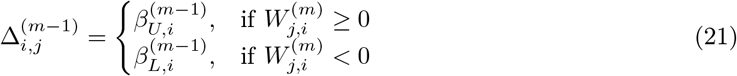

Plug (20) and (21) to (19), we get:

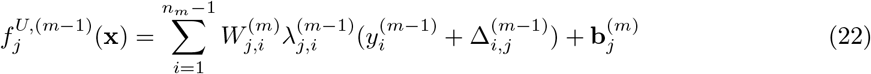

Imposing the relationship 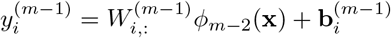, we obtain:

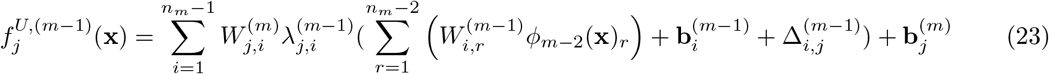

(23) can be simplified by defining 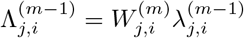:

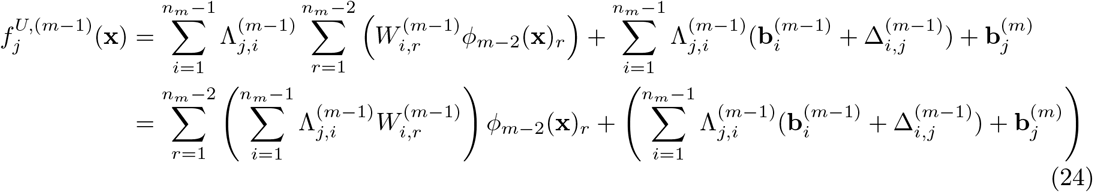

Note that 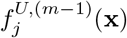 is not the bound that we desire. In reality, *ϕ*_*m*−2_(**x**) will also have “error”. As a result, we need to “propagate” this “error” to the input of the network where the error is known. To find the bound of the output of the network as a function of the bound of the input, we need to express *ϕ*_*m*−2_(**x**) = *σ*(**y**^(*m*−2)^) in f its upper bound, defined also by (13). Note that if we redefine a weight matrix 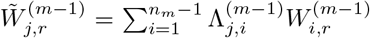, whether or not to use the upper or lower bound of *ϕ*_*m*−2_(**x**)_*i*_ will depend on the sign of 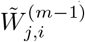. Consequently, we need to define:

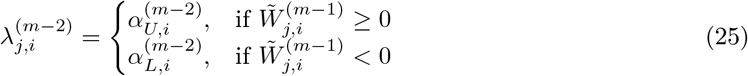

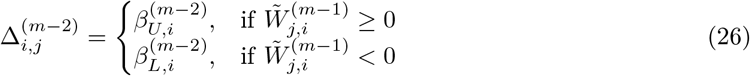

Plug (25), (26), and the definition of the upper and lower bound of *ϕ*_*m*−2_(**x**) to (24), we get:

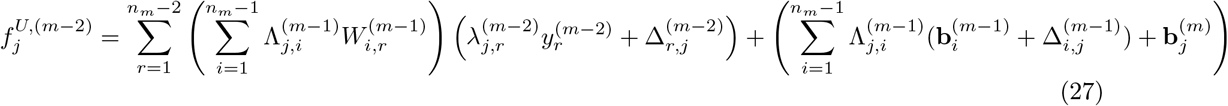

Again, impose the relationship 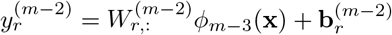

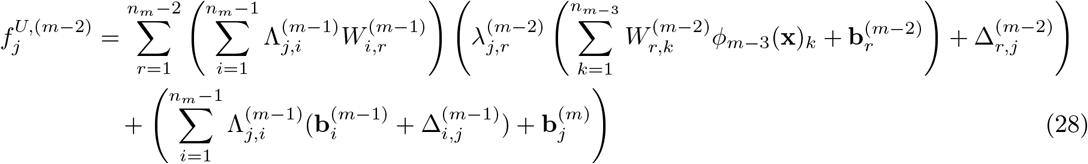

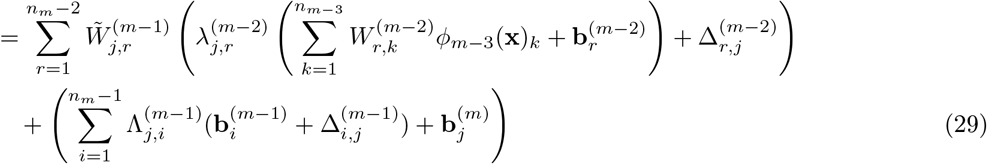

To simplify (29), define 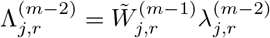, we get:

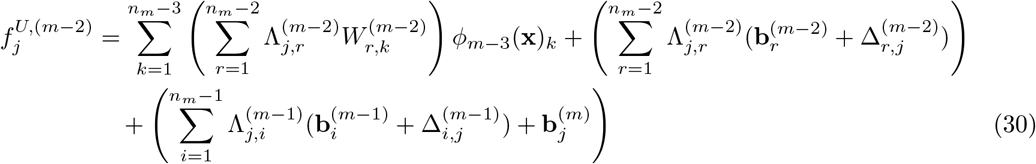

If we define:

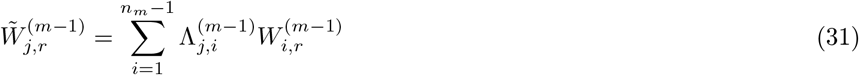

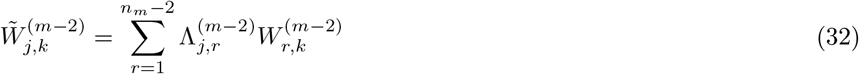

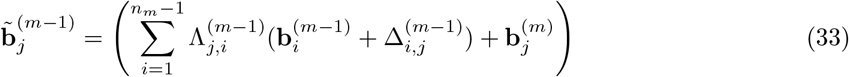

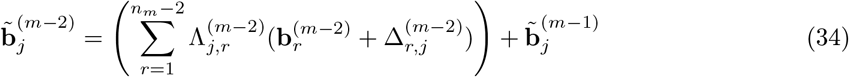

we can simplity (24) and (30) to:

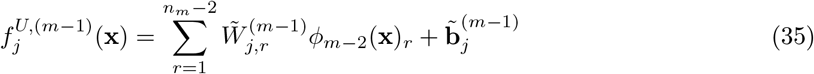

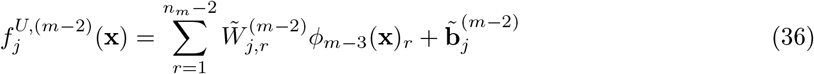

Note that from 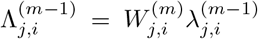 and 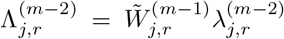, we get obtain an iterative relationship for Λ as:

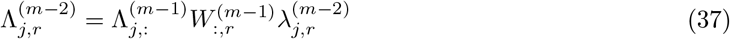

In the situation when *m* = 3, one can write out the general form of the upper bound as:

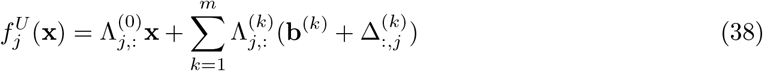

The result for deriving the lower bound is essentially the same as above and we will direct readers to the original paper.

When the input **x**_0_ can be perturbed within a ball of radius *ϵ*, **B**(**x**_0_, *ϵ*), we can write out its upper bound as:

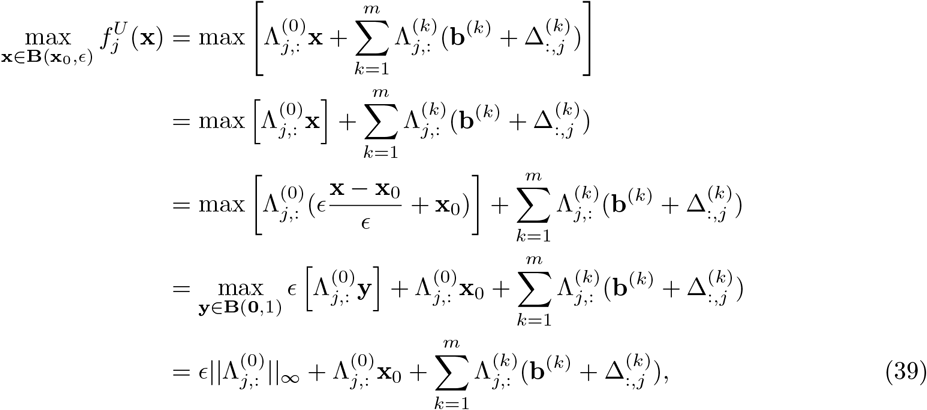

where 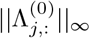 denotes the maximum absolute value of the vector 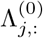

## References

[1] Kelly Street, Davide Risso, Russell B. Fletcher, Diya Das, John Ngai, Nir Yosef, Elizabeth Purdom, and Sandrine Dudoit. Slingshot: cell lineage and pseudotime inference for single-cell transcriptomics. BMC Genomics, 19(1):477, December 2018.

[2] Xiaojie Qiu, Qi Mao, Ying Tang, Li Wang, Raghav Chawla, Hannah A Pliner, and Cole Trapnell. Reversed graph embedding resolves complex single-cell trajectories. Nature Methods, 14(10):979–982, October 2017.

[3] Wouter Saelens, Robrecht Cannoodt, Helena Todorov, and Yvan Saeys. A comparison of single-cell trajectory inference methods. Nature Biotechnology, 37(5):547–554, May 2019.

[4] Daniel E. Wagner and Allon M. Klein. Lineage tracing meets single-cell omics: opportunities and challenges. Nature Reviews Genetics, 21(7):410–427, July 2020.

[5] Chang Su, Zichun Xu, Xinning Shan, Biao Cai, Hongyu Zhao, and Jingfei Zhang. Cell-type-specific co-expression inference from single cell RNA-sequencing data. Nature Communications, 14(1):4846, August 2023.

[6] Ziwei Wang, Hui Ding, and Quan Zou. Identifying cell types to interpret scRNA-seq data: how, why and more possibilities. Briefings in Functional Genomics, 19(4):286–291, July 2020.

[7] Megan Crow, Anirban Paul, Sara Ballouz, Z. Josh Huang, and Jesse Gillis. Characterizing the replicability of cell types defined by single cell RNA-sequencing data using MetaNeighbor. Nature Communications, 9(1):884, February 2018.

[8] Laurens van der Maaten and Geoffrey Hinton. Visualizing data using t-sne. Journal of Machine Learning Research, 9(86):2579–2605, 2008.

[9] Leland McInnes, John Healy, and James Melville. UMAP: Uniform Manifold Approximation and Projection for Dimension Reduction. 2018. Publisher: arXiv Version Number: 3.

[10] Jiarui Ding and Aviv Regev. Deep generative model embedding of single-cell RNA-Seq profiles on hyperspheres and hyperbolic spaces. Nature Communications, 12(1):2554, May 2021.

[11] Jiarui Ding, Anne Condon, and Sohrab P. Shah. Interpretable dimensionality reduction of single cell transcriptome data with deep generative models. Nature Communications, 9(1):2002, May 2018.

[12] Benjamin Szubert, Jennifer E. Cole, Claudia Monaco, and Ignat Drozdov. Structure-preserving visualisation of high dimensional single-cell datasets. Scientific Reports, 9(1):8914, June 2019.

[13] Tim Sainburg, Leland McInnes, and Timothy Q Gentner. Parametric UMAP embeddings for representation and semi-supervised learning. 2020. Publisher: arXiv Version Number: 4.

[14] Wei Dong, Charikar Moses, and Kai Li. Efficient k-nearest neighbor graph construction for generic similarity measures. In Proceedings of the 20th international conference on World wide web, pages 577–586, Hyderabad India, March 2011. ACM.

[15] Adam Paszke, Sam Gross, Francisco Massa, Adam Lerer, James Bradbury, Gregory Chanan, Trevor Killeen, Zeming Lin, Natalia Gimelshein, Luca Antiga, Alban Desmaison, Andreas Köpf, Edward Yang, Zach DeVito, Martin Raison, Alykhan Tejani, Sasank Chilamkurthy, Benoit Steiner, Lu Fang, Junjie Bai, and Soumith Chintala. PyTorch: An Imperative Style, High-Performance Deep Learning Library. Curran Associates Inc., Red Hook, NY, USA, 2019.

[16] Diederik P. Kingma and Jimmy Ba. Adam: A Method for Stochastic Optimization. 2014. Publisher: arXiv Version Number: 9.

[17] Marius Lange, Volker Bergen, Michal Klein, Manu Setty, Bernhard Reuter, Mostafa Bakhti, Heiko Lickert, Meshal Ansari, Janine Schniering, Herbert B. Schiller, Dana Pe’er, and Fabian J. Theis. CellRank for directed single-cell fate mapping. Nature Methods, 19(2):159–170, February 2022.

[18] Aimée Bastidas-Ponce, Sophie Tritschler, Leander Dony, Katharina Scheibner, Marta Tarquis-Medina, Ciro Salinno, Silvia Schirge, Ingo Burtscher, Anika Böttcher, Fabian Theis, Heiko Lickert, and Mostafa Bakhti. Massive single-cell mRNA profiling reveals a detailed roadmap for pancreatic endocrinogenesis. Development, page dev.173849, January 2019.

[19] Gioele La Manno, Ruslan Soldatov, Amit Zeisel, Emelie Braun, Hannah Hochgerner, Viktor Petukhov, Katja Lidschreiber, Maria E. Kastriti, Peter Lönnerberg, Alessandro Furlan, Jean Fan, Lars E. Borm, Zehua Liu, David Van Bruggen, Jimin Guo, Xiaoling He, Roger Barker, Erik Sundström, Gonçalo Castelo-Branco, Patrick Cramer, Igor Adameyko, Sten Linnarsson, and Peter V. Kharchenko. RNA velocity of single cells. Nature, 560(7719):494–498, August 2018.

[20] Hannah Hochgerner, Amit Zeisel, Peter Lönnerberg, and Sten Linnarsson. Conserved properties of dentate gyrus neurogenesis across postnatal development revealed by single-cell RNA sequencing. Nature Neuroscience, 21(2):290–299, February 2018.

[21] F. Alexander Wolf, Philipp Angerer, and Fabian J. Theis. SCANPY: large-scale single-cell gene expression data analysis. Genome Biology, 19(1):15, December 2018.

[22] Tim Stuart, Andrew Butler, Paul Hoffman, Christoph Hafemeister, Efthymia Papalexi, William M. Mauck, Yuhan Hao, Marlon Stoeckius, Peter Smibert, and Rahul Satija. Comprehensive Integration of Single-Cell Data. Cell, 177(7):1888–1902.e21, June 2019.

[23] Huan Zhang, Tsui-Wei Weng, Pin-Yu Chen, Cho-Jui Hsieh, and Luca Daniel. Efficient Neural Network Robustness Certification with General Activation Functions. 2018. Publisher: arXiv Version Number: 1.

[24] Kaidi Xu, Zhouxing Shi, Huan Zhang, Yihan Wang, Kai-Wei Chang, Minlie Huang, Bhavya Kailkhura, Xue Lin, and Cho-Jui Hsieh. Automatic perturbation analysis for scalable certified robustness and beyond. Advances in Neural Information Processing Systems, 33, 2020.

[25] Kaidi Xu, Huan Zhang, Shiqi Wang, Yihan Wang, Suman Jana, Xue Lin, and Cho-Jui Hsieh. Fast and Complete: Enabling complete neural network verification with rapid and massively parallel incomplete verifiers. In International Conference on Learning Representations, 2021.

[26] Shiqi Wang, Huan Zhang, Kaidi Xu, Xue Lin, Suman Jana, Cho-Jui Hsieh, and J Zico Kolter. Beta-CROWN: Efficient bound propagation with per-neuron split constraints for complete and incomplete neural network verification. arXiv preprint arXiv:2103.06624, 2021.

[27] G. Edward W. Marti, Steven Chu, and Stephen R. Quake. Aging causes changes in transcriptional noise across a diverse set of cell types. preprint, Bioinformatics, June 2022.

[28] Stephen J. Fleming, Mark D. Chaffin, Alessandro Arduini, Amer-Denis Akkad, Eric Banks, John C. Marioni, Anthony A. Philippakis, Patrick T. Ellinor, and Mehrtash Babadi. Unsupervised removal of systematic background noise from droplet-based single-cell experiments using CellBender. Nature Methods, 20(9):1323–1335, September 2023.

[29] Diederik P Kingma and Max Welling. Auto-Encoding Variational Bayes. 2013. Publisher: arXiv Version Number: 11.

